# Retrospective Computational and Immunoinformatics Validation of Clinically Evaluated *Necator americanus* Protein Vaccine Candidates: *Na*-APR-1, *Na*-GST-1, and *Na*-ASP-2

**DOI:** 10.64898/2026.09.02.748823

**Authors:** Yanick Aqua Stong Tangan, G Takop Nchanji, Marcel Nyuylam Moyeh, Teke Mary Efeti, Methodius Shinyuy Lahngong, Kwi Pilate Nkineh, Abey Blessings Ayuk, Junior Ekunidi Engarimbi, Che Roland Achungu, Cabirou Mounchili Shintouo, Dulandzev Melvis Berinyuy, Ntang Emmaculate Yaah, Bernis Neneyoh Yengo, Tamnjong Beltine Muyer, Ketura Yaje Gwei, Brinate Teke Tebo, Nfiaboh Gaston Nyah, Stephen Mbigha Ghogomu, Robert Adamu Shey

## Abstract

Hookworm disease, primarily caused by *Necator americanus*, affects about 472 million people worldwide and contributes substantially to global disease burden, yet no approved vaccine is currently available. The clinical failure of *Na*-ASP-2 protein due to IgE-mediated hypersensitivity highlights the need for safe, immunogenic hookworm vaccines and emphasizes the importance of rigorous pre-clinical safety screening. Using an integrated immunoinformatics approach, this work retrospectively evaluated the safety and immunogenicity of three clinical hookworm protein vaccine candidates: *Na*-APR-1, *Na*-GST-1, and *Na*-ASP-2. Toxigenicity and allergenicity predictions correctly identified *Na*-ASP-2 as toxigenic and allergenic, consistent with its documented clinical failure, while *Na*-APR-1 exhibited a favourable safety profile; *Na*-GST-1 showed inconsistent allergenicity signals warranting experimental validation. Comprehensive epitope prediction identified abundant CTL, HTL, B-cell, and cytokine-inducing epitopes across all candidates, with *Na*-APR-1 demonstrating the broadest epitope repertoire. HLA population coverage analysis indicated broad global applicability across endemic regions. Molecular docking with TLR4 revealed that all antigens interact with the receptor with binding energies more favourable than the positive control agonist, with *Na*-GST-1 and *Na*-APR-1 displaying the strongest predicted affinities. Normal mode analyses predicted stable antigen-TLR4 complex dynamics across all candidates. Immune simulations predicted robust, memory-driven humoral and cellular responses for *Na*-APR-1 and *Na*-ASP-2, while *Na*-GST-1 showed markedly attenuated simulated immunogenicity despite favourable structural and receptor-binding characteristics. These findings support continued clinical development of *Na*-APR-1, highlight unresolved immunogenic discordances for *Na*-GST-1 requiring experimental verification, and collectively provide a validated computational framework for advancing rational hookworm vaccine design.

**Author summary:** Hookworm infection affects hundreds of millions of people worldwide, yet no vaccine exists to prevent it. Three protein-based vaccine candidates have already been tested in clinical trials: one failed because it triggered allergic reactions, while the other two showed promise but raised unanswered questions. In this study, we used computer-based tools to compare all three candidates side by side, asking how well each one might be recognized by the immune system and whether any carried hidden safety risks. Interestingly, our predictions correctly flagged the candidate that failed in humans as risky, showing that this type of computational screening can predict potential problems before they reach clinical trials. One candidate stood out as the most balanced option, combining good safety with a strong predicted immune response. A second candidate demonstrated favorable performance in several analyses but yielded unexpectedly weak results in others, despite evidence of efficacy in vaccinated individuals, suggesting that these discrepancies may reflect limitations of the computational approaches rather than inherent shortcomings of the vaccine itself. Our findings support using these computational approaches as an early, low-cost step to help researchers prioritize the most promising hookworm vaccine candidates before committing to expensive lab and clinical testing.

## Introduction

Hookworm disease, caused predominantly by *Necator americanus* (*N. americanus*), is among the most consequential neglected tropical diseases (NTDs). Globally, it affects an estimated 472 million people and disproportionately burdens impoverished communities in sub-Saharan Africa, Southeast Asia, and Latin America [1, 2]. *N. americanus* infects humans via skin penetration by L3 larvae, which subsequently migrate to the small intestine. The parasite feeds on blood, causing iron-deficiency anemia, impaired child development, adverse pregnancy outcomes, and about 4.1 million DALYs lost annually, with an economic burden exceeding $139 billion [3].

Control currently relies on mass drug administration (MDA) with benzimidazole anthelmintics, but its effectiveness is limited by rapid reinfection, lack of lasting immunity, and emerging drug resistance with reduced efficacy [4]. These shortcomings underscore the urgent need for alternative measures, including a prophylactic vaccine capable of conferring durable, memory-mediated protection, as recognized in the WHO 2030 NTD roadmap [5]. Modeling studies on the economic and epidemiological impact of an effective hookworm vaccine have shown that, when implemented in high-transmission settings, it would be both highly cost-effective and a potential cost-saving biotechnology [6]. In addition to providing direct protection, a vaccine would help reduce the emergence of anthelmintic drug resistance while also decreasing the likelihood of disease recurrence [7].

To date, no licensed vaccine is available for the prevention of human hookworm infection [8]. However, considerable research and development efforts targeting *N. americanus* have largely focused on three leading vaccine candidates, *N. americanus* Ancylostoma-secreted protein 2 (*Na*-ASP-2) [9], *N. americanus* Aspartic Proteinase-1 (*Na*-APR-1) [10], and *N. americanus* Glutathione S-transferase-1 (*Na*-GST-1) [11]. *Na*-ASP-2 was selected for its role in the transition to parasitism [12] and its ability to elicit a protective Th2-type immune response [9]. However, the Phase I clinical trial was halted after participants experienced urticarial reactions, linked to pre-existing *Na*-ASP-2-specific IgE antibodies that triggered type I hypersensitivity responses [13]. *Na*-APR-1 and *Na*-GST-1 are crucial for adult hookworm survival, as they facilitate the degradation and utilization of host hemoglobin obtained through blood feeding [10, 11]. *Na*-APR-1 facilitates hemoglobin digestion, whereas *Na*-GST-1 detoxifies heme released during this process. Both antigens have demonstrated strong protective immunity against hookworm infection in vaccinated animals [14] and have subsequently undergone testing for human vaccination in clinical trials involving adults and children [14]. Vaccination in adults was deemed safe and effective in eliciting specific IgG responses; therefore, Phase II clinical trials are currently underway [15]. The co-administration of recombinant *Na*-APR-1(M74) and *Na*-GST-1 was well tolerated in school-aged Gabonese children and elicited significant IgG antibody responses [16].

The development of an effective vaccine against *N. americanus* remains a global health priority [3]. Although two vaccine candidates that combine *Na*-APR-1 and *Na*-GST-1 have advanced to clinical evaluation due to their crucial roles in triggering immune responses and supporting parasite survival [15], several critical questions remain unanswered by experimental data alone. Clinical trials assess antigens under different conditions, limiting direct comparison, and the molecular and epitope basis of *Na*-ASP-2-associated IgE hypersensitivity remains unclear. Computational immunoinformatics provides a standardized framework for comparing vaccine antigens, identifying epitope-level immune mechanisms, and generating testable hypotheses regarding antigen immunogenicity and safety. Such analyses are particularly valuable for understanding why highly immunogenic candidates, such as *Na*-ASP-2, progressed successfully through preclinical evaluation yet failed during clinical testing because of IgE-mediated hypersensitivity. This highlights the essential need for rigorous preclinical safety profiling, especially in populations with prior exposure to the parasite.

To our knowledge, these antigens are the most extensively characterized *N. americanus* vaccine candidates evaluated in both preclinical and clinical studies, making them appropriate benchmark antigens for assessing the performance of computational immunoinformatics tools. To support the hookworm vaccine research community in addressing these challenges, we performed homology modeling, HLA allele-specific epitope prediction, and immune simulation analyses on three well-characterized hookworm vaccine candidates: *Na*-APR-1, *Na*-GST-1, and *Na*-ASP-2, to systematically compare their predicted capacities to elicit humoral and cellular immune responses. This comparison is not intended as definitive validation but rather as a preliminary evaluation of the extent to which *in silico* predictions are consistent with existing experimental and clinical evidence. Such benchmarking provides an initial assessment of the extent to which computational immunoinformatics can support the prioritization of vaccine candidates before resource-intensive experimental and clinical evaluation. Although these findings cannot substitute for experimental validation, they contribute to the growing body of evidence supporting the use of computational immunoinformatics in early-stage hookworm vaccine candidate selection.

## Methodology

### Protein sequence and structure retrieval

The full-length amino acid sequences of *Na*-APR-1, *Na*-GST-1, and *Na*-ASP-2 (UniProtKB accession numbers Q9N9H3, D3U1A5, and Q7Z1H1, respectively) were obatined from the [UniProtKB](https://www.uniprot.org/blast) database in FASTA format and saved as text files. The three-dimensional structure of the human TLR4 and the TLR4 agonist, 50S ribosomal protein ribosomal protein L7/L12 (Locus RL7_MYCTU, UniProt ID: P9WHE3) from *Mycobacterium tuberculosis* was retrieved from the [Protein Data Bank](https://www.rcsb.org/) (PDB) and [UniProtKB](https://www.uniprot.org/blast) database respectively. Because it activates TLR4 signaling, the protein was used as a reference agonist and molecular control for docking and molecular dynamics simulations [17]. Although TLR4 is also involved in the induction of Th2- mediated immune responses [18], evidence indicates that it can interact with antigens from several helminths, including *Strongyloides stercoralis* and *O. volvulus* [18, 19].

### Research design

This study employed a three-phase immunoinformatics framework to comprehensively assess the safety, immunogenicity, and immune recognition potential of the vaccine candidates from multiple immunological perspectives.

### Epitope Characterization

Different prediction tools employing distinct algorithms, parameters, and experimentally validated datasets were used to identify diverse, immunologically relevant epitopes representing key vaccine attributes of each antigen. The assessment focused on: Safety-related properties; Potential immunomodulatory or immune-evasive (immune camouflage) characteristics; Capacity to induce antibody-mediated (humoral) immune responses; Ability to stimulate cellular immune responses; and potential to trigger cytokine-mediated responses that coordinate and bridge humoral and cellular immunity.

### Innate Immune Recognition Assessment

Structural bioinformatics approaches were used to predict and evaluate the interactions between each vaccine candidate and (TLR4), a key receptor involved in innate immune recognition and activation. This analysis aimed to determine the capacity of the candidates to effectively engage innate immune signaling pathways.

### *In Silico* Immune Response Simulation

Computational immune simulations were conducted for all vaccine candidates to model the dynamics of both primary and secondary immune responses over time. The simulations evaluated: Antibody-mediated (humoral) immunity; Cell-mediated immune responses; and Cytokine-driven immune responses, providing an integrated view of the overall immunological profile of each candidate.

### Signal peptide, sequence localization, and transmembrane domain prediction

The selected proteins were subjected to a series of preliminary analyses. Signal peptide prediction was conducted using the [SignalP 6.0](https://services.healthtech.dtu.dk/services/SignalP-6.0/) and [TOPCONS](https://topcons.cbr.su.se/) servers to distinguish classically secreted proteins from non-secretory proteins. Subcellular localization was assessed using [DeepLoc 2.1](https://services.healthtech.dtu.dk/services/DeepLoc-2.1/) and [WoLF PSORT](https://wolfpsort.hgc.jp/), while transmembrane regions were predicted using [DeepTMHMM 1.0](https://services.healthtech.dtu.dk/services/DeepTMHMM-1.0/) and [TOPCONS](https://topcons.cbr.su.se/) servers.

### Antigenicity, toxicity and allergenicity prediction

The antigenic potential of the vaccine candidates was evaluated using two alignment-independent prediction platforms, [VaxiJen v2.0](https://www.ddg-pharmfac.net/vaxijen/VaxiJen/VaxiJen.html) and [ANTIGENpro](http://scratch.proteomics.ics.uci.edu/). VaxiJen v2.0 predicts antigenicity by applying auto-and cross-covariance (ACC) transformation to protein sequences, converting them into uniform vectors based on key physicochemical properties of amino acids [20]. In contrast, ANTIGENpro utilizes protein microarray-derived antigenicity data and machine learning-based models to predict antigenic proteins, achieving an estimated cross-validation accuracy of approximately 76% [21]. To assess safety, the toxicity of each antigen was predicted using the ToxinPred2 server. The platform generates overlapping peptides and uses machine-learning models, including ET, ANN-LSTM, and hybrid ET+MERCI and DL+MERCI approaches, to predict potentially toxic regions [22]. The allergenic potential of the vaccine candidates was examined using [AllerTOP v2.0](https://www.ddg-pharmfac.net/allertop_v2/) and [AllergenFP v1.1](https://ddg-pharmfac.net/AllergenFP/). AllerTOP v2.0 uses ACC-based sequence transformation and machine-learning classifiers to distinguish allergenic from non-allergenic proteins, with reported accuracies exceeding 85% across diverse datasets [23]. Conversely, AllergenFP v1.1 predicts allergenicity using a descriptor-based binary fingerprinting approach that characterizes proteins according to their physicochemical properties [24].

### Protein conservation analyses

To evaluate potential cross-reactivity between the vaccine candidates and the human proteome, BLASTp searches were performed against the UniProtKB reviewed human proteome database. The amino acid sequences of *Na*-APR-1, *Na*-GST-1, and *Na*-ASP-2 were used as query sequences. The analysis aimed to identify identical sequence stretches shared between the vaccine candidates and human proteins. Previous studies have indicated that pathogen-derived peptides exhibiting sequence similarity to human proteins may bind the same HLA alleles, potentially promoting immune tolerance and reducing immunogenicity [25]. The BLAST search was conducted using customized parameters, including an E-value threshold of 0.01, the BLOSUM62 substitution matrix, filtering of low-complexity regions, and exclusion of gapped alignments. The E-value indicates the probability of a sequence match occurring by chance, with lower values reflecting more significant similarity [26]. While BLOSUM62, assigns scores based on observed amino acid substitutions, facilitating the identification of biologically meaningful sequence alignments [27].

### Physicochemical properties and solubility prediction

The physicochemical characteristics of the three vaccine candidates were evaluated using the [ProtParam](https://web.expasy.org/protparam/) web server. Parameters assessed included molecular weight (kDa), amino acid composition, theoretical isoelectric point (pI), estimated in vitro and in vivo half-life, instability index, aliphatic index, and the grand average of hydropathicity (GRAVY) [28]. Protein solubility was subsequently predicted using the NetSolP-1.0 and DeepSoluE servers. NetSolP-1.0 leverages deep-learning protein language models to predict solubility, providing state-of-the-art performance and improved generalizability across diverse datasets [29]. In contrast, DeepSoluE uses an LSTM-based neural network integrating physicochemical and amino acid representation features to predict recombinant protein solubility in bacterial expression systems [30].

### Prediction of linear B-cell epitopes

B-cell epitopes are antigenic determinants recognized by the immune system and represent the specific piece of the antigen to which B lymphocytes bind [31], and are vital in vaccine design. Linear B-cell epitopes were predicted using the [ABCpred](https://webs.iiitd.edu.in/raghava/abcpred/) server, which employs artificial neural networks for linear B-cell epitope prediction with a threshold set at 0.70 [32].

### Cytotoxic T lymphocytes (CTL) and Helper T-cell (HTL) epitope prediction

Cytotoxic T-lymphocyte (CTL) epitopes were predicted for the selected antigens using the publicly available [NetCTL 1.2](https://services.healthtech.dtu.dk/services/NetCTL-1.2/) server. This platform integrates three key components of antigen processing and presentation, namely MHC class I peptide binding, proteasomal C-terminal cleavage, and transporter associated with antigen processing (TAP) transport efficiency, to identify potential CTL epitopes. Although NetCTL 1.2 supports predictions across 12 MHC class I supertypes, only the A2, A3, and B7 supertypes were included in this study because together they provide population coverage of up to 90% [33]. Epitope selection was performed using the default prediction threshold score of 0.75. To identify HTL epitopes, the [NetMHCII 2.3](https://services.healthtech.dtu.dk/services/NetMHCII-2.3/) server was used to predict 15-mer peptides capable of binding human MHC class II molecules. The server employs artificial neural network-based algorithms to predict peptide interactions with HLA-DR, HLA-DQ, and HLA-DP alleles [34]. Predictions were generated using the default criteria for strong binders (SB) and weak binders (WB).

### Prediction of cytokine-inducing epitopes (IFN-γ, IL-2 IL-4, IL5, IL-17, IL-10 and TNF-α)

Interferon-gamma (IFN-γ) plays a pivotal role in both innate and adaptive immunity and has been recognized as a key indicator of protective Th1 responses against *Necator americanus* infection [35]. To identify IFN-γ-inducing regions, 15-mer epitopes from the selected antigens were predicted using the [IFNepitope](http://crdd.osdd.net/raghava/ifnepitope/) server. Interleukin-2 (IL-2), another cytokine associated with Th1- mediated protection against *N. americanus* [36], was evaluated by predicting 10-mer IL-2- inducing epitopes using the IL2Pred server [37]. Interleukin-4 (IL-4) is a hallmark cytokine of T-helper 2 (Th2) immune responses and is predominantly produced by CD4+ T cells during helminth infections [36]. The prediction of 15-mer IL-4-inducing epitopes for all vaccine candidates was carried out using the IL4Pred server [38]. Interleukin-5 (IL-5) has also been implicated in protective immunity against *N. americanus* [35]. Therefore, potential 15-mer IL-5-inducing epitopes were identified using the IL5Pred server, applying the default prediction threshold of 0.2 [39]. Interleukin-6 (IL-6) is a multifunctional cytokine involved in regulating inflammatory and immune responses during infection and tissue damage [40]. IL-6-inducing peptides within the\ vaccine candidates were predicted using the IL6Pred server [41]. Hookworm infection is associated with elevated constitutive and antigen-specific IL-10, which suppresses cellular immune responses [42]. To assess this immunomodulatory potential, 15-mer IL-10-inducing epitopes were predicted using the Random Forest-based model implemented in the IL10Pred server [43]. Finally, tumor necrosis factor-alpha (TNF-α) has been identified in murine studies as a soluble immune mediator potentially associated with vaccine-induced protection [35]. TNF-α-inducing epitopes within the vaccine candidates were predicted using the [TNFepitope](https://webs.iiitd.edu.in/raghava/tnfepitope/) server, employing the default prediction threshold value of 0.45 [44].

### Population coverage analysis

The [IEDB](http://tools.iedb.org/population/) Population Coverage tool was used to estimate the global coverage of CTL and HTL epitopes, including respective HLA alleles, across various geographical regions spanning multiple continents and sub-regions. This tool estimates population coverage for immune responses by analyzing HLA genotypic frequencies from the Allele Frequency database. It accommodates both class I and class II T cell epitopes, offering options for separate or combined coverage [45]. For this study, our focus was on the A2, A3, and B7 for MHC class I epitopes, and HLA-DR, HLA-DQ, and HLA-DP alleles for MHC class II epitopes, as these offer optimal population coverage for *N. americanus* endemic zones [46].

### Immune simulation

To further evaluate the immunogenic potential and immune response dynamics of the selected antigens, *in silico* immune simulations were performed using the C-ImmSim server [7]. C-ImmSim is an agent-based immune simulation platform that combines position-specific scoring matrix (PSSM)-based epitope prediction with machine learning algorithms to model complex immune interactions. The system simultaneously simulates three major mammalian immune compartments: (i) the bone marrow, where hematopoietic stem cells generate lymphoid and myeloid cell populations; (ii) the thymus, where naïve T cells undergo selection processes to prevent autoimmunity; and (iii) a tertiary lymphoid organ, such as a lymph node, where immune responses are initiated and regulated [47]. Immune simulations were conducted using the server’s default settings. Antigen administrations were scheduled at time steps 0, 84, and 168 to mimic successive immunizations. In the C-ImmSim framework, each time step corresponds to 8 hours, with the first injection occurring at time step 1 (time = 0) [48]. This approach enabled the modeling of both primary and secondary immune responses and the assessment of long-term immunological memory elicited by the vaccine candidates.

### Secondary structure and intrinsic disorder prediction

The secondary structural features of the vaccine candidates were predicted using the [PSIPREDv4.0](https://bioinf.cs.ucl.ac.uk/psipred/) and [RaptorX](http://raptorx.uchicago.edu/) Property web servers, while intrinsically disordered regions were identified using the [AIUPred](https://aiupred.elte.hu/) server. PSIPRED uses PSI-BLAST-derived PSSMs to predict protein secondary structure, achieving a Q3 accuracy of 81.6% under stringent cross-validation [49]. To complement these analyses, secondary structure prediction was also performed using the RaptorX Property server, which utilizes Deep Convolutional Neural Fields (DeepCNF), a deep learning framework capable of simultaneously predicting secondary structure elements, solvent accessibility, and intrinsically disordered regions. RaptorX Property has demonstrated high predictive accuracy for both protein secondary structure and disorder characterization [50]. Disordered proteins constitute an important class of antigens in numerous human pathogens and are frequently associated with protective immune responses [51]. AIUPred combines a biophysics-based prediction framework with deep learning techniques to accurately identify intrinsically disordered protein regions [52].

### Tertiary structure prediction, refinement and validation

The 3D structural models of the vaccine candidates were predicted using [ColabFold](https://support.microsoft.com/en-us/internet-explorer/this-website-doesn-t-work-in-internet-explorer), a user- friendly platform that implements AlphaFold2 technology within the Google Colab environment [53]. ColabFold integrates MMseqs2 with AlphaFold2 or RoseTTAFold to enable rapid and accurate prediction of protein structures and complexes [54]. The predicted 3D models were subsequently refined using the [GalaxyRefine](http://alaxy.seoklab.org/) server. GalaxyRefine enhances protein model quality by rebuilding and repacking side chains and applying molecular dynamics simulations, improving local structural accuracy and overall model relaxation [55]. Following refinement, the structural integrity and overall quality of the vaccine candidate models were evaluated using the [ProSA-web](https://prosa.services.came.sbg.ac.at/prosa.php) server. ProSA-web is widely utilized for the identification of structural errors in experimentally determined protein structures, theoretical models, and engineered proteins by assessing their overall quality against known protein structures [56]. Further validation was carried out using the [ERRAT](https://saves.mbi.ucla.edu/) server, which assesses the reliability of protein models by analyzing non-bonded atom-atom interaction patterns and comparing them with those observed in high-resolution crystallographic structures. This provides an additional measure of structural accuracy and model quality [57].

### Prediction of discontinuous B-lymphocyte (DBL) epitopes

It has been estimated that >90% of B-cell epitopes are discontinuous (conformational) [58]. To predict these discontinuous B-cell epitopes for the validated 3D structures produced as described in the structure prediction section below, the [ElliPro](http://tools.iedb.org/ellipro/) tool was used. The ElliPro tool integrates three algorithms: approximating the protein shape as an ellipsoid, calculating the residue protrusion index (PI), and clustering neighboring residues based on their PI values [59].

### Binding pocket prediction molecular docking, binding affinity, and interaction analyses of vaccine candidates with the TLR4 receptor

The [CASTp 3.0](https://cfold.bme.uic.edu/castpfold/) server was employed to predict potential binding pockets in the chimeric vaccine construct. The server uses an alpha-shape algorithm to identify and characterize protein cavities and pockets based on their geometric and topological properties [60]. To evaluate the ability of the vaccine candidates to engage the innate immune system through TLR4, protein-protein docking analyses were performed using the [HawkDock](http://cadd.zju.edu.cn/hawkdock/) server. The 3D structures of each vaccine candidate, the TLR4 agonist, and the TLR4 receptor were submitted to HawkDock as ligand and receptor molecules, respectively. HawkDock is an integrated platform that combines molecular docking with molecular mechanics/generalized Born surface area (MM/GBSA) calculations to assess protein-protein interactions and estimate binding free energies [61]. Among the generated docking poses, the complex exhibiting the lowest MM/GBSA binding free energy was selected for subsequent analyses, as lower energy values are generally indicative of more stable, native-like, and thermodynamically favorable interactions [62]. The resulting complexes were visualized using PyMOL. Also, binding affinities between the vaccine candidates, the TLR4 agonist, and the TLR4 receptor were estimated using the [PROtein binDIng enerGY](https://wenmr.science.uu.nl/prodigy/manual) (PRODIGY) server, which predicts binding free energy (ΔG) based on intermolecular contact features [63]. In parallel, the docked complexes were examined using [PDBsum](http://www.ebi.ac.uk/thornton-srv/databases/pdbsum/Generate.html) server, which generates 2D schematic representations of molecular interactions at the protein-protein interface from 3D structural coordinates. These complementary analyses were chosen because HawkDock provides MM/GBSA-based binding free energy estimations, whereas PRODIGY evaluates binding strength using contact-based energetic parameters, thereby offering independent and complementary insights into complex stability and interaction quality [63].

### Normal mode analyses

Identification of flexible regions within protein structures is crucial for understanding their biological functions and dynamic behaviour [64]. To evaluate the structural flexibility and stability of the docked antigen-TLR4 complexes, normal mode analysis (NMA) was performed using the **[iMODS](https://imods.chaconlab.org/)** server. This analysis provides insights into the internal dihedral coordinates of protein complexes and characterizes their collective functional motions. The essential dynamics simulation module of iMODS was employed to investigate the conformational behaviour, energy minimization, structural stability, and atomic mobility of the vaccine candidate-TLR4 complexes. The server predicts the most likely motions and deformation patterns of the complexes using a range of structural and dynamical parameters, including B-factors, root mean square deviation (RMSD), eigenvalues, deformability profiles, covariance matrices, and elastic network models [65]. These parameters collectively provide valuable information on the flexibility, motion, and stability of the docked complexes, thereby facilitating the assessment of their potential biological relevance.

## Results

### Study design

This study provides a retrospective computational validation of three hookworm vaccine candidates with established preclinical and clinical profiles. The study was designed firstly to assess various epitopes that in each vaccine candidate represent responsiveness to vaccine candidates’ potential adverse side reactions, immunomodulation (camouflage) property, antibody-based (humoral) responses, cellular immune responses, as well as cytokine-based responses (regulation and mediation between humoral and cellular responses). Different computational tools with a unique set of parameters and individually trained on distinct experimental datasets were used to achieve this. Secondly, through structural simulations, the study predicts the ability of each vaccine candidate to be recognized by the innate immune TLR4 receptor. Lastly, immune simulation was also performed to model the primary and secondary humoral, cellular, and cytokine-mediated responses over time.

### Signal peptide, sequence localization, and transmembrane domain prediction

Both servers predicted the presence of signal peptides only in *Na*-APR-1, while no signal peptides were predicted for either *Na*-GST-1 or *Na*-ASP-2. For TM domain prediction, all three proteins were predicted by both DeepTMHMM and TOPCONS servers to contain no transmembrane domains. Also, both the DeepLocPro-1.0 and WoLF PSORT servers predicted *Na*-ASP-2 to be localized in the extracellular space. On the other hand, *Na*-GST-1 was predicted to be localized in the cytoplasm by both servers, while *Na*-APR-1 was predicted to be localized in the Lysosome/Vacuole and Plasma membrane by both servers, respectively (**Table 1**).

**Table 1:**
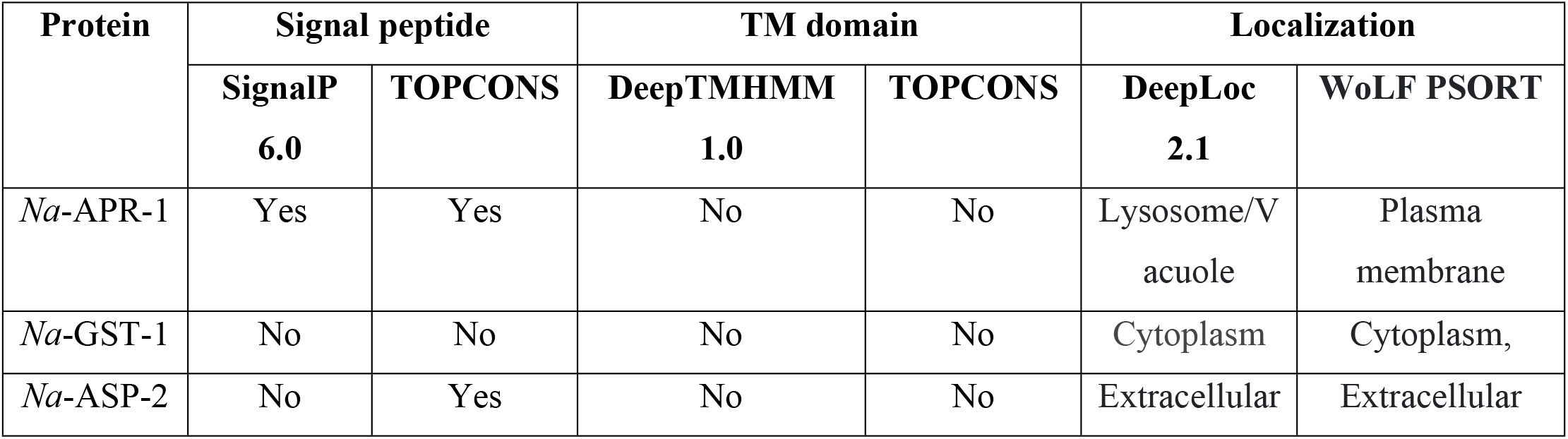
Signal peptide, sequence localization, and transmembrane domain prediction.

### Antigenicity, toxigenicity, and allergenicity predictions

The antigenicity predicted for *Na*-APR-1, *Na*-GST-1, and *Na*-ASP-2 showed scores of 0.43, 0.55, and 0.80, respectively, on VaxiJen v2.0, and *Na*-APR-1 and *Na*-ASP-2 showed 0.59 and 0.92, on ANTIGENpro. While *Na*-GST-1 showed a score of 0.30 on ANTIGENpro. The threshold values for both servers were set at the default (0.5). Therefore, according to VaxiJen v2.0 and ANTIGENpro scores, all antigens were predicted to be significantly antigenic except for *Na*-GST-1, which had a lower antigenicity score as predicted by ANTIGENpro (**Table 2**).

**Table 2.** Vaccine-desired properties prediction.

| Vaccine candidate | Antigenicity |  | Solubility |  | Allergenicity |  | Usability | Toxicity |
| --- | --- | --- | --- | --- | --- | --- | --- | --- |
|  | ANTIGENpro | VaxJen | DeepSoluE | NetSolP-1.0 | AllerTOP | AllergenFP | NetSolP-1.0 | ToxinPred2 |
| *Na*-APR-1 | 0.59 | 0.43 | 0.7614 | 0.4358 | Non-allergen | Non-allergen | 0.3787 | Non-toxin |
| *Na*-GST-1 | 0.30 | 0.55 | 0.5258 | 0.5806 | Non-allergen | Allergen | 0.4958 | Non-toxin |
| *Na*-ASP-2 | 0.92 | 0.80 | 0.7724 | 0.5965 | Allergen | Allergen | 0.4298 | Toxin |

To ensure the safety of the vaccine candidates, toxicity and allergenicity levels were predicted for each antigen. Toxic proteins or peptides have been reported to induce dementia (e.g., Alzheimer’s disease) and heart disease in some patient groups [66]. *Na*-APR-1 and *Na*-GST-1 were predicted to have no toxic peptides, while *Na*-ASP-2 was predicted to contain toxic peptides (**Table 2**).

The AllerTOP v.2 and AllergenFP servers showed that *Na*-APR-1 was non-allergenic, while *Na*-ASP-2 was predicted to contain allergenic epitopes by both servers. *Na*-GST-1, on the other hand, was predicted to contain allergenic epitopes by AllergenFP but was predicted to be non-allergenic by AllerTOP v.2.

### Physicochemical properties and solubility analysis

Analysis of the physicochemical properties of the antigens supports their suitability for use in hookworm vaccination strategies. Overall, the vaccine candidates show acidic-to-basic predicted isoelectric points (pI 5.74 to 8.20) and favourable expression half-lives across systems (yeast >20 h, *E. coli* >10 h, and mammalian reticulocytes ∼30 h). Most candidates exhibit acceptable protein stability based on instability index values, with GRAVY scores remaining negative (−0.122 to - 0.389), indicating improved water interaction/solubility [67]. Aliphatic indices were high (59.10-88.01), suggesting thermostable proteins [68], while molecular weights ranged from ∼23.7 to ∼72.6 kDa (**Table 3**).

**Table 3.** Physicochemical properties of *N. americanus* vaccine antigens.

| Vaccine candidate | theoretical isoelectric point (pI) | *in vivo* half-life | *in vitro* half-life | instability index | aliphatic index | molecular weight (MW) | grand average of hydropathicity |
| --- | --- | --- | --- | --- | --- | --- | --- |
| *Na*-APR-1 | 6.67 | >20 h (yeast, *in vivo*)<br>>10 h (*E. coli*, *in vivo*) | 30 h (mammalian reticulocytes, *in vitro*) | 52.11 | 79.39 | 49554.78 | -0.122 |
| *Na*-GST-1 | 5.74 | >20 h (yeast, *in vivo*)<br>>10 h (*E. coli*, *in vivo*) | 30 h (mammalian reticulocytes, *in vitro*) | 30.76 | 88.01 | 23679.38 | -0.147 |
| *Na*-ASP-2 | 8.20 | >20 h (yeast, *in vivo*)<br>>10 h (*E. coli*, *in vivo*) | 30 h (mammalian reticulocytes, *in vitro*) | 35.39 | 59.10 | 22605.91 | -0.389 |

The solubility scores were predicted to be 0.4358, 0.5806, and 0.5965 for *Na*-APR-1, *Na*-GST-1, and *Na*-ASP-2, respectively, by the NetSolP 1.0 server, but 0.7614, 0.5258, and 0.7724, respectively, on the DeepSoluE server. The protein exhibited varying solubility predictions (0.4358 and 0.7724). Furthermore, the NetSolP 1.0 server predicted all the antigens to have a usability score ranging from 0.3787 to 0.4958 (**Table 2**).

### Protein conservation analyses

A significant degree of conservation among all antigens was noted in the human proteome (ranging from 22.8% to 53%) based on a BLAST search conducted against the UniProtKB database (**Table 4**). For *Na*-APR-1, the server predicted 50 homologs ranging from 22.8% to 52.5% identity. Also, 27 homologs were predicted for *Na*-GST-1, ranging from 23.9% to 36%, whereas *Na*-ASP-2 showed 38 homologs, ranging from 23.5% to 44.3%.

**Table 4.** Conservation analysis shows identical sequence stretches common to some human proteins.

| Vaccine candidate | Rank | Homolog (*Homo sapiens* (Human)) | Protein name | % Identity |
| --- | --- | --- | --- | --- |
| *Na*-APR-1 (50 hits) | 1 | P07339 | Cathepsin D | 52.5% |
|  | 2 | O96009 | Napsin-A | 50.3% |
|  | 3 | P14091 | Cathepsin E | 43.1% |
|  | 4 | P0DJD7 | Pepsin A-4 | 42.1% |
| *Na*-GST-1 (27 hits) | 1 | O60760 | Hematopoietic prostaglandin D synthase | 35.6% |
|  | 2 | P21266 | Glutathione S-transferase Mu 3 | 29.7% |
|  | 3 | Q7RTV2 | Glutathione S-transferase A5 | 27.7% |
|  | 4 | P09211 | Glutathione S-transferase P | 28.9% |
| *Na*-ASP-2 (38 hits) | 1 | P54108 | Cysteine-rich secretory protein 3 | 34% |
|  | 2 | P16562 | Cysteine-rich secretory protein 2 | 33.7% |
|  | 3 | Q6UWM5 | GLIPR1-like protein 1 | 33.3% |
|  | 4 | Q6UXB8 | Peptidase inhibitor 16 | 32.8% |

### Linear and discontinuous B-cell epitope prediction

The ABCpred server predicted 20 linear B-cell epitopes for *Na*-APR1; 12 for *Na*-ASP-2; and 9 for *Na*-GST-1 (**Table 5**). In contrast, for discontinuous (conformational) B-cell epitopes, a total of 15, 8, and 8 residues were predicted to be found in 11, 6, and 4 different discontinuous epitopes for *Na*-APR-1, *Na*-GST-1, and *Na*-ASP-2, respectively, with scores ranging from 0.50 to 0.988 (**Table 5** and **S1 Table 1**). Overall, *Na*-APR-1 predicted substantially more linear and continuous B-cell epitopes, followed by *Na*-GST-1, which was higher than *Na*-ASP-1. All antigens contained multiple high-scoring sequences, indicating strong potential for antibody recognition and serving as markers for humoral response evaluation.

**Table 5.** Abundance of different epitope types in the vaccine candidates.

| Vaccine candidate | CTL | HTL | IL-4 | IL-5 | IL-2 | IFN-γ | IL-6 | IL-10 | TNF-α | LBL | DBL |
| --- | --- | --- | --- | --- | --- | --- | --- | --- | --- | --- | --- |
| *Na-APR-1* | 45 | 503 | 259 | 142 | 234 | 73 | 35 | 93 | 75 | 20 | 11 |
| *Na-GST-1* | 13 | 53 | 136 | 116 | 199 | 15 | 28 | 58 | 27 | 12 | 6 |
| *Na-ASP-2* | 15 | 312 | 121 | 74 | 190 | 28 | 10 | 56 | 32 | 09 | 4 |

### Helper T (CD4+) and Cytotoxic T lymphocyte (CD8+) epitope prediction

The NetCTL 1.2 server predictions indicate that *Na*-APR-1, *Na*-GST-1, and *Na*-ASP-2, respectively, contain 45, 13, and 15 CTL epitopes (approximately 9-mers), with most specific to the HLA-A3 super-type. The NetMHCII 2.3 server predictions of the 15-mer overlapping HTL epitopes for all antigens (targeting HLA-DR, HLA-DQ, and HLA-DP alleles) revealed strong binders (SB) or weak binders (WB) according to their binding affinity. For all the antigens, the SB epitopes showed binding affinity values ranging from 1.3 nM to 1.3 mM; a total of 868 SB were predicted (HLA-DR: 272, HLA-DQ: 256, HLA-DP: 341). *Na*-APR-1, *Na*-GST-1, and *Na*-ASP-2 yielded (HLA-DR: 111, HLA-DQ: 114, HLA-DP: 278), (HLA-DR: 27, HLA-DQ: 17, HLA-DP: 9), and (HLA-DR: 134, HLA-DQ: 125, HLA-DP: 53), respectively (**Table 5**).

### Prediction of cytokine-inducing epitopes (IFN-γ, IL-2 IL-4, IL5, IL-17, IL-10 and TNF-α)

IL4pred server predicted *Na*-APR-1, *Na*-GST-1, and *Na*-ASP-2 to contain >100 IL-4-inducing epitopes (**Table 5**). These abundant IL-4-inducing epitopes suggest that all proteins can stimulate the desired Th2 responses required for protection against *N. americanus* in the human host. The IL2pred webserver predicted *Na*-APR-1, *Na*-GST-1, and *Na*-ASP-2 to contain >180 IL-2-inducing epitopes (**Table 5**). For IL-5, 142, 116, and 74 inducing epitopes were predicted to be present in *Na*-APR-1, *Na*-GST-1, and *Na*-ASP-2, respectively (**Table 5**). The IL-10pred server predicted 93 epitopes for *Na*-APR-1, 62 epitopes for *Na*-GST-1, and 58 epitopes for *Na*-ASP-2. Also, IL-6 server predicted 35, 28, and 10 IL-6-inducing epitopes present in *Na*-APR-1, *Na*-GST-1, and *Na*-ASP-2, respectively (**Table 5**). Furthermore, a total of 73, 15, and 28 potential IFN-γ-inducing epitopes (15-mers) were predicted for *Na*-APR-1, *Na*-GST-1 and *Na*-ASP-2, respectively. Finally, for TNF-α, 75, 27, and 32 potential epitopes were predicted to be present in *Na*-APR-1, *Na*-GST-1, and *Na*-ASP-2, respectively (**Table 5 and S1 Table 2**).

### Population coverage analysis

The IEDB population coverage calculation tool predicted that the overall coverage for MHC class I CTL epitopes was 63.96% for all the antigens, while MHC class II HTL epitopes showed a coverage of 100% for all antigens. For specific regions, the predicted coverage for MHC class I CTL epitopes was: Central Africa (47.97%), East Africa (47.75%), South Africa (41.61%), West Africa (47.99%), Southeast Asia (47.34%), East Asia (51.98%) and South America (66.11%) for all the antigens (**S1 Fig. 1**). The highest percentage coverage for MHC class I, was found in South America at 66.11%, while Southeast Asia recorded the lowest with 47.34%. Similarly, for MHC class II HTL epitopes, the coverage was predicted to be 100% in all selected regions for all the antigens except for the South African population which recorded a 51.56% coverage (**S1 Fig. 2**).

**Fig. 1.**
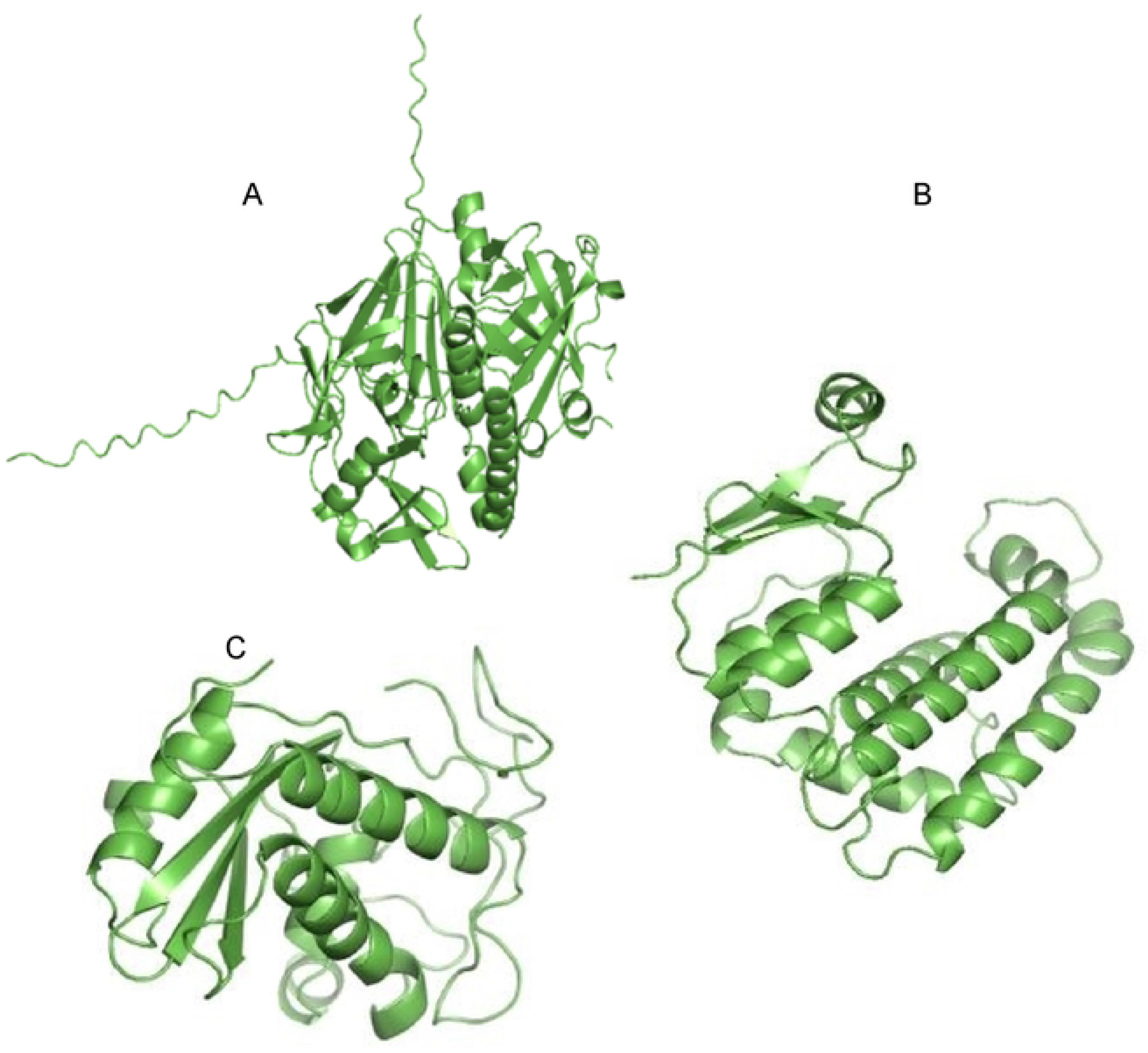
**Predicted 3D structures of N. americanus antigens generated using AlphaFold 2**. Panels A-C correspond to (A), Na-APR-1 (B), Na-GST-1 (C), and Na-ASP-2. Most residues are predicted with either very high (pLDDT >90) confidence for Na-GST-1 and Na-ASP-2 or high (70-90) confidence for Na-APR-1 and the chimeric antigens.

**Fig 2.**
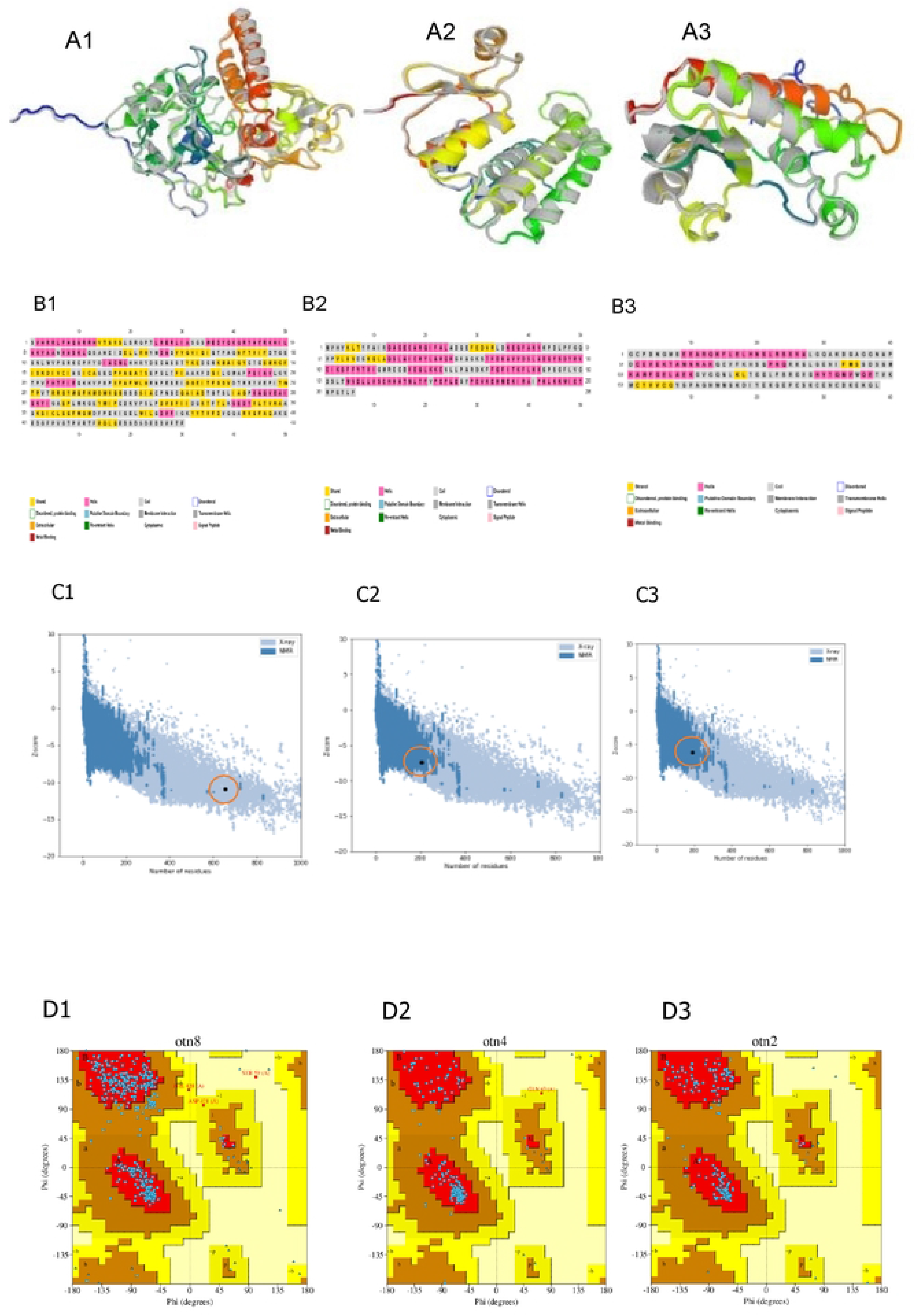
Structural modelling and quality assessments of vaccine candidates. A1) Na-APR-1, A2) Na-GST-1, and A3) Na-ASP-2 refined models. B1) Na-APR-1, B2) Na-GST-1, and Na-ASP-2 secondary structural features. C1) Na-APR- 1, C2) Na-GST-1, and C3) Na-ASP-2 graphical presentation of high-accuracy predictions relative to experimentally solved structures. D1) Na-APR-1, D2) Na-GST-1, and D3) Na-ASP-2. Ramachandran analyses reveal most residues in predicted models are localized in favoured regions.

### Secondary structure and disorder prediction, tertiary structure modelling, refinement, and validation

Secondary structure and disorder prediction analysis showed that *Na*-APR-1, *Na*-GST-1, and *Na*- ASP-2 are comprised of approximately 13%, 67.9%, and 26.7% alpha helices, 29.3%, 7.8%, and 5.8% beta-strands, and 57.8%, 24.3%, and 67.9% coils, respectively (**Fig. 2B 1-B3)**. Solvent accessibility analysis revealed that all 3 antigens contained 40%, 35%, and 45%, respectively, of residues fully exposed, 27%, 30%, and 21%, respectively, were partially exposed, and 31%, 33%, and 32%, respectively, were buried (**Table 6**). The AIUPred web server predicted ∼15% and ∼85% of residues were in the ordered and disordered regions, respectively for *Na*-APR-1. Likewise, for *Na*-ASP-2, ∼48% and ∼52% of residues were in the ordered and disordered regions, respectively. For *Na*-GST-1, ∼100% of residues were predicted to be in the ordered region and none in the disordered region (**S1 Fig. 6**).

**Table 6.** Secondary structure features of the *N. americanus* antigens.

| Antigen | Secondary structure |  |  |  |  |  |
| --- | --- | --- | --- | --- | --- | --- |
| | α-helices (%) | β-strands (%) | coiled coils (%) | Exposed (%) | Partially exposed (%) | Buried (%) |
| *Na*-APR-1 | 13 | 29.3 | 57.8 | 40 | 27 | 31 |
| *Na*-GST-1 | 67.9 | 7.8 | 24.3 | 35 | 30 | 33 |
| *Na*-ASP-2 | 26.7 | 5.8 | 67.9 | 45 | 21 | 32 |

The 3D structure of the antigens predicted using AlphaFold2 on the ColabFold interface (**Fig 1**) yielded 5 models for each antigen after refinement. Among these 5 models, “models 2, 1, and 2”, were selected for *Na*-APR-1, *Na*-GST-1, and *Na*-ASP-2 were selected with parameters GDT-HA (0.9756), (0.9951), and (0.9908), RMSD (0.349), (0.267) and (0.273), MolProbity (1.548), (1.445) and (1.660), and Rama favoured (98.8), (99.0) and (98.4) respectively. Models with the best quality scores: RMSD <4 Å, MolProbity score <2, and Ramachandran plot score > 97% were then selected for further analysis (**Fig. 2A 1-A3**).

ProSA-web revealed Z-scores of −8.89, −7.3, and −6.23 for *Na*-APR-1, *Na*-GST-1, and *Na*-ASP-2 (**Fig. 2C1-C3**), indicating high accuracy of prediction relative to experimentally solved structures on proteins in the PDB database. Ramachandran plot analysis (**Fig. 2D1-D3**) of the modelled proteins revealed 87.8 %, 94.5%, and 93.3% of residues for *Na*-APR-1, *Na*-GST-1, and *Na*-ASP- 2, respectively. Furthermore, the ERRAT server revealed an overall structural quality factor of 94.94%, 97.98%, and 95.05% for *Na*-APR-1, *Na*-GST-1, and *Na*-ASP-2, respectively. From a structural perspective, all candidates exhibited good structural integrity as predicted, as shown by the ERRAT server.

### Binding pocket prediction molecular docking, binding affinity, and interaction analyses of vaccine candidates with the TLR4 receptor

Molecular docking was performed to evaluate the interaction between the refined chimeric vaccine structure and TLR4 following prediction of its protein-protein interaction pocket (**S1 Fig 2**). HawkDock server predicted the binding interaction between the refined 3D structure of the antigens and TLR4. One hundred models were generated, visualized, and selected based on their ranking from the HawkDock server, with model 1 selected for downstream analysis and compared with the control antigen (**Fig 3, A-C and S1 Fig 1**). For *Na*-APR-1-TLR4, *Na*-GST-1-TLR4, and *Na*-ASP-2-TLR4, the binding energies were −291.11, −310.47, and −244.47, respectively, while for agonist-TLR4, the binding energy was −215.66. Overall, all antigens exhibited more favorable binding energies to TLR4 than the control antigen (−215.66), suggesting stronger predicted binding and potentially more favorable receptor-ligand interactions (**Table 7**).

**Fig 3.**
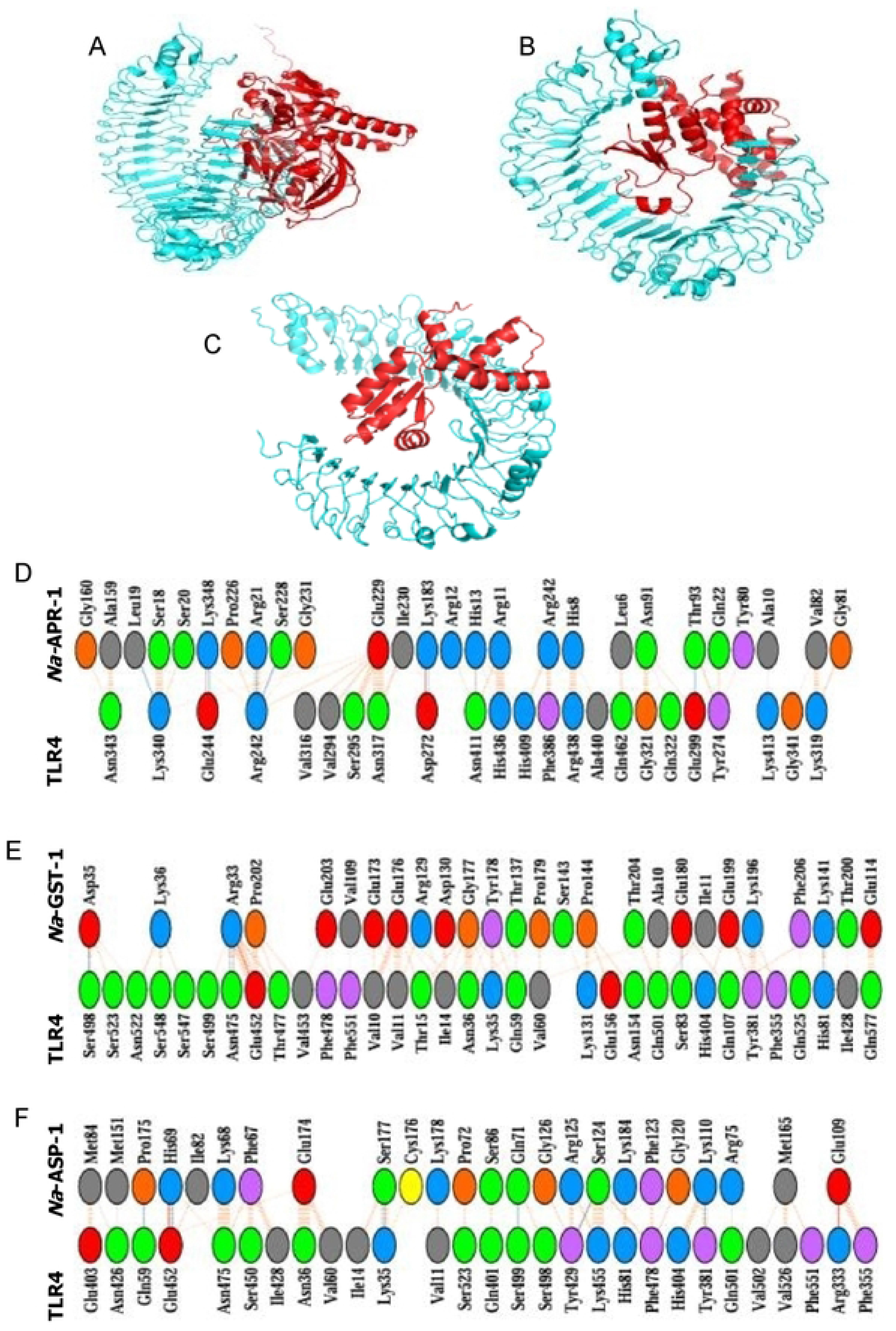
**Molecular docking protein-protein interaction pocket prediction and of the vaccine candidates with the TLR4 receptor were conducted**. A) Na-APR-1-TLR4 complex, B) Na-GST-1-TLR4 complex, and C) Na-FUS-1-TLR4 complex. The docked complex of the vaccine candidates (red) and the Toll-like receptor 4 chain (cyan) is shown. D) Na-APR-1-TLR4 complex, E) Na-GST-1-TLR4, and F) Na-FUS-1-TLR4 complex. The interaction network between the vaccine candidates and TLR4 is illustrated, with hydrogen bonding interactions depicted in blue. Non-bonded interactions and salt bridges are represented in orange. Charged residues are color-coded: positive in blue, negative in red, neutral in green, aliphatic in grey, and aromatic in purple.

**Table 7.**
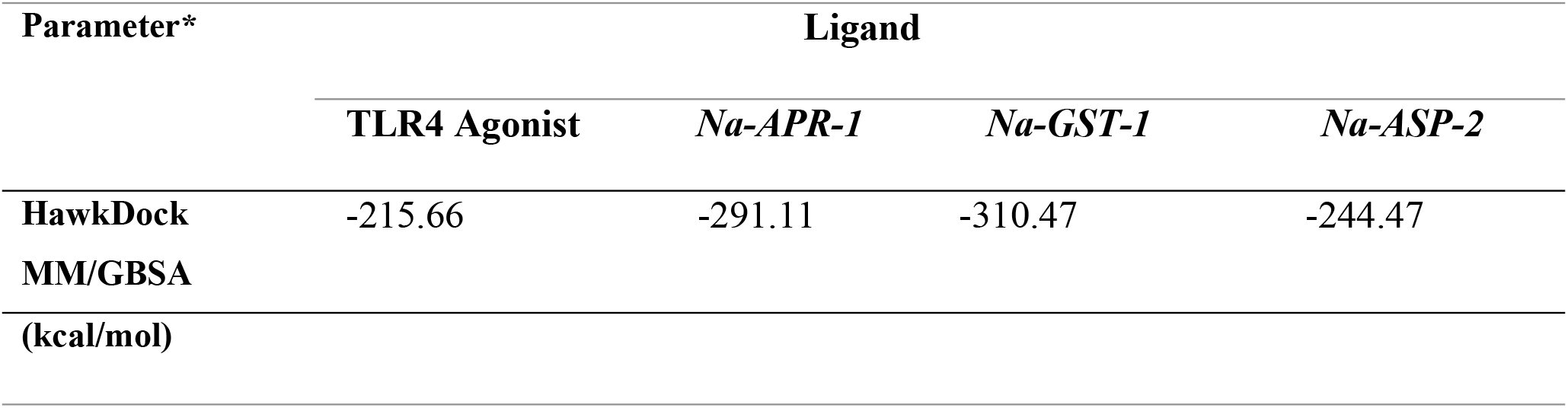
Energy characterization of the top-ranked vaccine candidate-TLR4 docked clusters.

To further investigate antigen-receptor interactions, the selected model of the antigens-TLR4 complex was assessed for interface interactions using PDBsum and binding affinity using the PrODIGY server. The PDBsum server predicted the formation of 7, 5, and 6 hydrogen bonds, 2, 1, 2, and salt bridges, and 229, 201, 168, and non-bonded contacts between 23, 33, and 28 residues from the TLR4 and 26, 26, and 24 residues from *Na*-APR-1, *Na*-GST-1, and *Na*-ASP-2, respectively. For the agonist-TLR4 complex, the server predicted 1 hydrogen bond, 5 salt bridges, and 94 non-bonded contacts between 20 residues from TLR4 and 16 residues of the agonist (**Fig 3D-F and S1 Fig 1**). Overall, the antigens demonstrated favorable interactions with TLR4 compared to the agonist with TLR4. The antigen-TLR4 complexes showed relative binding free energies (ΔG) of −13.3 Kcal/mol, −16.2 Kcal/mol, and −14.0 Kcal/mol (for *Na*-APR-1-, *Na-*GST- 1-, and *Na-*ASP-2-TLR4), respectively (**S1 Table 3**). While for the agonist-TLR4 complex, the binding energy was −11.5 Kcal/mol. The corresponding predicted Kd values for the antigens-TLR4 were 1.6e-10, 1.4e-12, and 5e-11, respectively, while the agonist-TLR4 complex had a Kd value of 3.4e-9, indicating stronger predicted binding of the antigens to TLR4 compared to the agonist (**S1 Table 3**).

### Normal mode analyses

The deformability graph showed distinct peak regions, indicating where the main-chain residues of the antigens-TLR4 complex undergo the greatest deformation. The hinge regions identified correspond to areas of high deformability (**Fig. 4A 1-3**). The B-factor plot shows the relationship between NMA mobility and the antigens-TLR4 complex, indicating the average RMSD values of the docked complex (**Fig 4B1-3**). B-factor values indicate the uncertainty associated with each atom. The computed eigenvalues for the antigens-TLR4 complexes were 2.5863241e-07, 8.145360e-05, and 2.239416e-05, which reflect motion stiffness with respect to each normal mode (**Fig. 4C 1-3**). The eigenvalue estimates the energy needed to deform the structure. Each normal mode of the complex is depicted by individual (purple) and cumulative (green) variance in the variance bar. It is worth noting that there is a negative correlation between the variance and the eigenvalue (**Fig. 4D. 1-3**). Furthermore, the interactions between the antigens and TLR4 in the complex are depicted using a covariance matrix. The related motions among various pairs of residues are indicated by correlated (red), uncorrelated (white), and anti-correlated (blue) atomic movements within the antigens-TLR4 complex (**Fig. 4E 1-3**). An elastic network map was also created to show atom pairs linked by springs in the antigens-TLR4 complex. Each dot in the graph denotes a spring connecting the corresponding pair of atoms. The color of the dots indicates their stiffness; darker shades of grey represent stiffer regions, while lighter dots signify more flexible areas (**Fig. 4F 1-3**). All the findings from the normal mode analyses suggest the favourable interaction and stability within the antigen-TLR4 complex.

**Fig 4.**
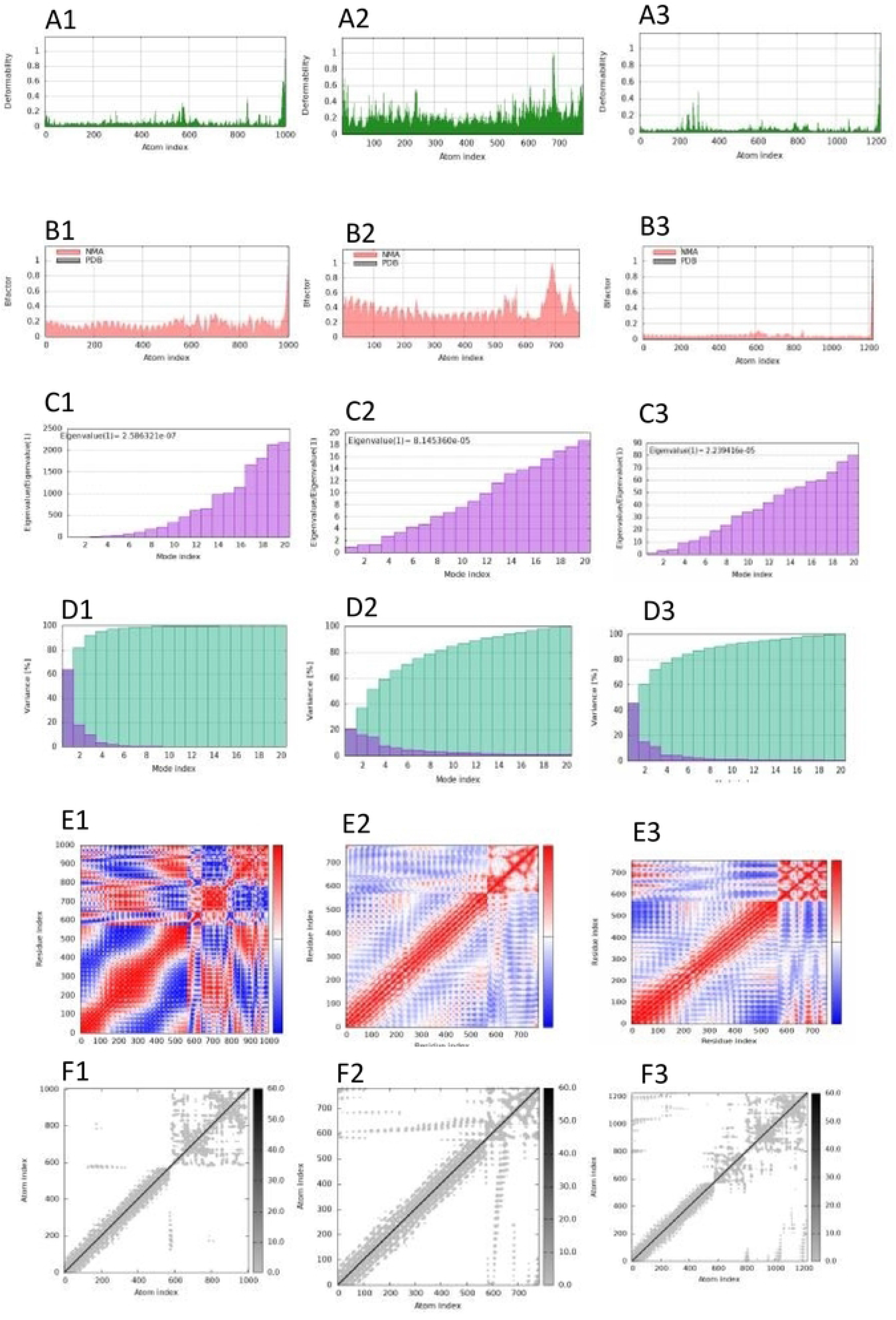
Molecular dynamics simulation of the vaccine candidate-TLR4 docked complex. (A1-3) Display the main chain deformability of the various antigen-TLR4 complexes. (B1-3) The B-factor quantifies the uncertainty associated with each atom. (C1-3) The eigenvalue indicates the motion stiffness related to each normal mode. (D1-3) The variance map shows individual (red) and cumulative (green) variances. (E1-3) The covariance graph illustrates the mobility of residue pairs, indicating correlated (red), uncorrelated (white), or anti-correlated (blue) movements. (F1-3) The elastic network model represents pairs of atoms connected by springs, with darker greys indicating greater stiffness of the springs: 1 = Na-APR-1-TLR4 complex, 2= Na-GST-1-TLR4 complex, and 3= Na-ASP2-1-TLR4 complex).

### Immunization simulation

To comparatively evaluate the immunogenic potential of individual *N. americanus* antigens *in silico* simulations with the C-ImmSim server, the same three-dose immunization schedule was performed as previously reported [69]. The C-ImmSim server predictions revealed consistent and robust immune responses with distinct patterns in the primary, secondary, and tertiary response categories (**Fig 5).** During the primary response (**Fig. 5A-C**), all antigens (except *Na*-GST-1) induced high levels of IgM antibodies, reflecting initial immune activation. The secondary and tertiary responses resulted in a marked rise in IgG1 and IgG2 (IgG1+IgG2), along with increased IgM and IgG+IgM antibody levels. A gradual reduction in antigen levels accompanied this. *Na*-GST-1 antigens did not induce either primary or secondary immune responses (**Fig 5B**). Immune memory was supported by the sustained elevation of IFN-γ levels and the increased T-helper cell population (Th2 response) across the exposure period (**Fig. 5D-5F**).

**Fig 5.**
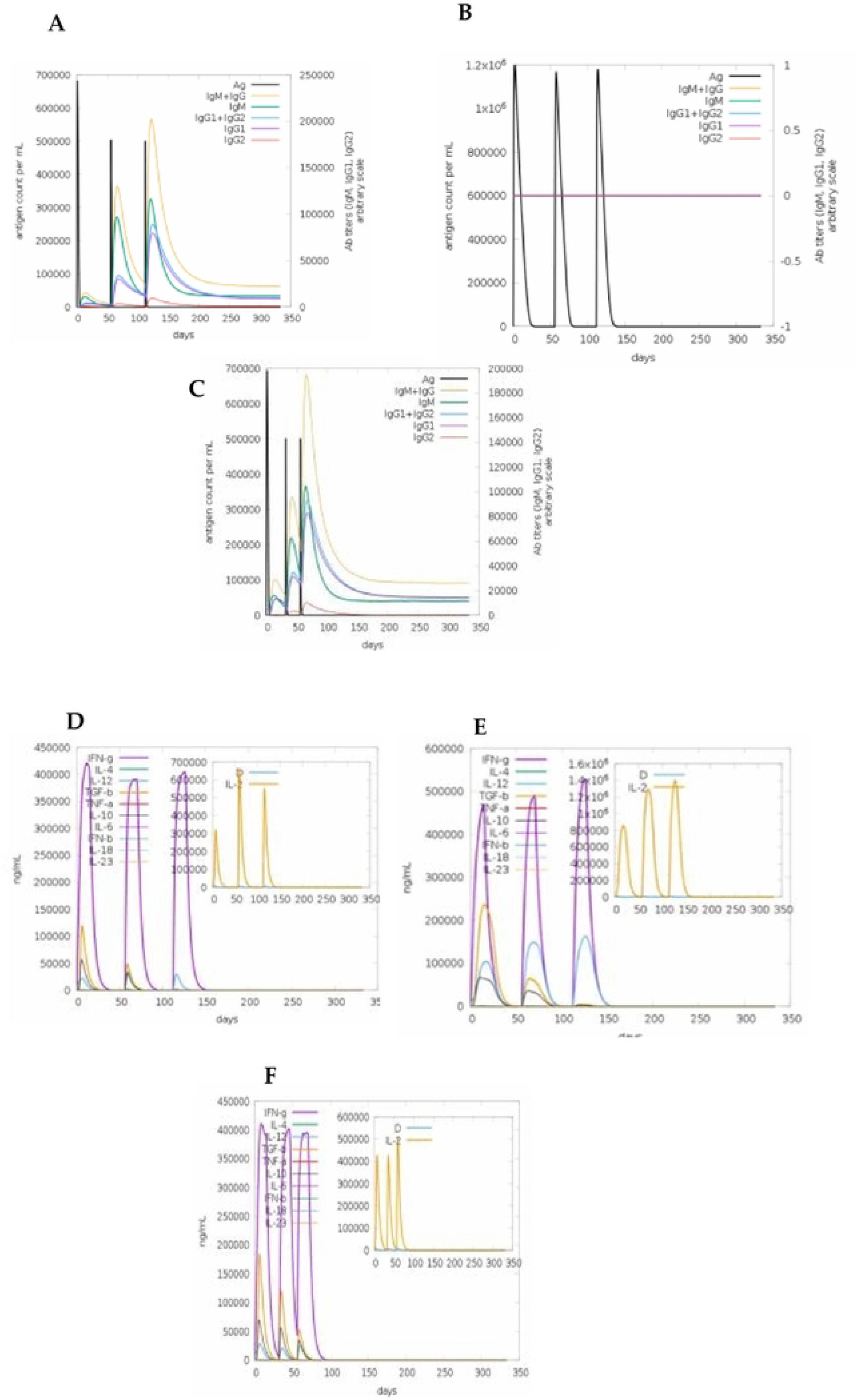
Virtual immune response of antibodies and cytokine responses for both vaccine candidates. Panels A) Na-APR-1, B) Na-GST-1, and C) Na-ASP-2 indicate virtual immune responses for both antigens. Immunoglobulin production in response to antigen injections (black vertical lines); specific subclasses are indicated as coloured peaks. Panels D) Na-APR-1, E) Na-GST-1, and F) Na-ASP-2 show virtual cytokine profiles for both candidate antigens with three injections given 4 weeks apart. The larger plot shows cytokine levels after the injections. The insert plot shows the IL-2 level with the Simpson index, D, indicated by the dotted line. D is a measure of diversity. An increase in D over time indicates the emergence of different epitope-specific dominant clones of T-cells. The smaller the D value, the lower the diversity.

**Fig 6.**
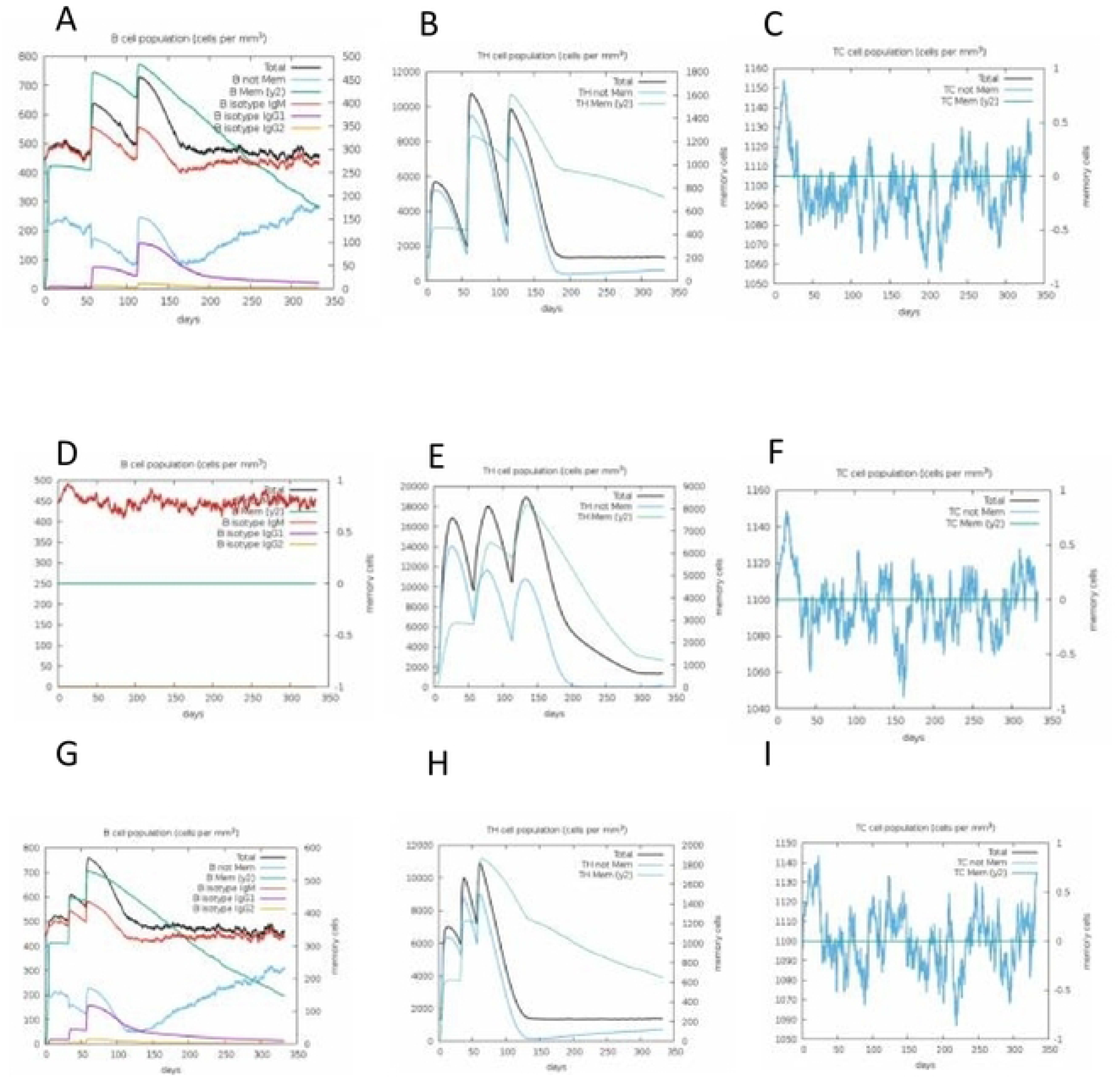
**Immune simulation of B- and T-cell responses for both vaccine candidates**. Upper and middle panels indicate A,D) B-cell, B,E) T Helper cell, and C, F) T cytotoxic cell responses for Na-APR-1 and Na-GST-1; while the lower panels are representations of G) B-cell, H) T Helper cell, and I) T cytotoxic cell responses for Na-ASP-2.

Furthermore, the simulations indicate (**Fig 6**) a high response in B-cell populations to all antigens (except *Na*-GST-1), as well as both TH (helper) and TC (cytotoxic) cell populations, further supporting the development of immune memory for both antigens. Overall, all antigens elicited comparable immune-cell responses. For all antigens except *Na*-GST-1 (**Fig 6D**), the total B-cell populations (**Fig 6A, 6G)** plateaued at roughly 340 cells/mm³. In contrast, the total T-helper (TH) cell populations for all the antigens declined sharply by about day 100, reaching approximately 200 memory TH cells/mm³ (**Fig. 6B, 6E, and 6H**). Also, the concentration of T cytotoxic cell population was almost constant across 350 days from the point of antigen inoculation (**Fig 6C, 6F, and 6I**). Overall, the simulations predicted slightly higher levels of antibodies against *Na*-ASP-2 compared to *Na*-APR-1, which was predicted to be higher than *Na*-GST-1.

## Discussion

This study is a retrospective computational validation of hookworm vaccine candidates with known clinical profiles [9, 15, 70]. Using an integrated immunoinformatics pipeline, we assessed the immunogenicity and safety of vaccine candidates through epitope prediction, safety profiling, structural modelling, molecular docking, and immune simulation. The findings were consistent with experimentally reported outcomes, providing mechanistic insights into antigen behaviour and supporting the use of *in-silico* approaches as valuable tools for early-stage vaccine screening and development [15, 16].

The objective of this study was to predict the immunogenicity and other important vaccine-related parameters of the separate antigens using immunoinformatics. *Na*-APR-1 and *Na*-GST-1 were predicted to be non-toxic, while *Na*-ASP-2 was identified as containing toxic peptides. Interestingly, *Na*-ASP-2 was predicted to be allergenic, a finding that is consistent with its discontinued Phase I trial due to IgE-mediated reactions [13]. The concordance between computational predictions and observed clinical outcomes highlights the need for a thorough immunoinformatics screening as an effective early warning tool in vaccine development. The consistent identification of *Na*-ASP-2 as allergenic by two independent prediction platforms further supports the reliability of this finding and its value in preventing adverse outcomes during clinical trials. In contrast, *Na*-APR-1 was predicted to be non-allergenic by both servers, consistent with its acceptable safety profile observed in Phase I and II clinical trials [70]. Notably, *Na*-GST- 1 presented a discordant result: while the server classified it as non-allergenic, the other predicted allergenic potential. The inconsistency may reflect limitations of the computational tools used rather than an inherent deficiency of the vaccine itself. Alternatively, it may be related to the similarity of *Na*-GST-1 to human GST proteins, which could induce cross-reactive immune responses or immune tolerance. Such sequence conservation with host proteins may trigger cross-reactive immune responses or promote immune tolerance, phenomena well-documented for GST- family antigens [71]. Although data from clinical studies confirm *Na*-GST-1 can elicit specific IgG responses without significant adverse events in vaccinated adults [15], the allergenic prediction requires rigorous experimental validation. *Na*-APR-1 showed the greatest similarity to human proteins, particularly Cathepsin D (up to 52.5% identity), suggesting potential immune evasion through molecular mimicry [72] (**Table 4**). Despite this, *Na*-APR-1 has demonstrated robust immunogenicity in clinical settings [70], suggesting that surface-exposed non-conserved epitopes rather than the conserved catalytic core may be the primary immunogenic targets. These findings underscore the importance of epitope-level filtering to exclude host-homologous regions in rational vaccine refinement. Assessment of humoral and cellular epitope content revealed that all three antigens harbour substantial predicted B-cell, CTL, and HTL epitopes, supporting their capacity to engage both MHC class I and class II antigen presentation pathways (**Table 2**). Among the individual antigens, *Na*-APR-1 yielded the highest number of linear B-cell epitopes and predicted HTL epitopes, while *Na*-ASP-2 was most antigenic by both servers. *Na*-GST-1, despite its structural richness, had the lowest antigenicity score on ANTIGENpro (0.30), a finding consistent with its high human proteome similarity that may promote immunological tolerance through shared HLA epitope presentation [25]. For CTL epitopes, it was predicted that the majority of the epitopes were from the HLA-A3 super-type, which, together with HLA-A2 and HLA-B7, can provide up to 90% population coverage in endemic regions [33].

Cytokine-inducing epitope analysis demonstrated that all antigens are predicted to stimulate a broad spectrum of immune mediators relevant to anti-hookworm immunity. All candidates harboured substantial IL-4- and IL-5-inducing epitopes, consistent with the Th2-skewed immune environment characteristic of protective responses against helminths [36]. *Na*-APR-1 yielded the highest counts of IFN-γ and TNF-α-inducing epitopes, supporting the capacity for mixed Th1/Th2 responses considered important for durable vaccine-mediated protection [73]. All antigens were predicted to induce an IL-10 response, warranting attention since IL-10 is known to maintain an immunosuppressed milieu during active hookworm infection [42], and a high IL-10 epitope density could potentially dampen vaccine-induced effector responses. Population coverage analysis confirmed broad global applicability, with predicted MHC class I CTL coverage of 63.96% across endemic regions and 100% MHC class II HTL coverage for all antigens.

To evaluate the capacity of each antigen to engage the innate immune system, molecular docking with the TLR4 receptor was performed with an *M. tuberculosis* 50S ribosomal protein as a positive control agonist. All three vaccine candidates exhibited more favourable binding energies than the control agonist. Notably, *Na*-GST-1 demonstrated the highest predicted TLR4 binding affinity among the three candidates. NMA revealed that all antigen-TLR4 complexes were structurally stable, with *Na*-GST-1 and *Na*-ASP-2 displaying higher structural rigidity (larger eigenvalues) and *Na*-APR-1 exhibiting greater conformational flexibility. This flexibility may allow *Na*-APR-1 to accommodate diverse TLR4 conformational states, though it may also reduce binding residence time [64].

The immune simulation results suggest that *Na*-ASP-2 exhibited a more sustained primary immune response and more specific and long-lasting secondary and tertiary immune responses, followed by *Na*-APR-1, as reflected by higher and sustained levels of IgM and IgG1+IgG2 antibodies. Both antigens show promise in triggering a mix of Th1 andTh2responses, which is essential for targeting hookworm through vaccination. *Na*-GST-1, on the other hand, failed to induce either primary or secondary humoral immune responses in the simulation, with negligible IgM or IgG production and absent B-cell expansion. This is the most significant internal contradiction of the present study, given that Na-GST-1 demonstrated the strongest predicted TLR4 binding affinity among all three antigens. We propose two complementary mechanistic explanations. First, the extensive sequence homology between *Na*-GST-1 and the human proteome may promote immune tolerance to *Na*- GST-1-derived epitopes. Secondly, the C-ImmSim platform may systematically underpredict immunogenicity for antigens whose epitope repertoire closely resembles host proteins, as the simulation model is trained on known human pathogen epitopes and may not adequately represent tolerance mechanisms for self-similar sequences [47]. Importantly, clinical data demonstrate that *Na*-GST-1 does elicit specific IgG responses in vaccinated adults [15], confirming that the simulation failure reflects a known model limitation for this antigen class rather than an absence of true immunogenic potential. The findings indicate that vaccine immunogenicity cannot be adequately assessed using a single computational measure; instead, a multi-parameter integrative approach combining safety, structural, binding, and immune response analyses offers the most reliable framework for prioritizing candidates. Overall, *Na*-APR-1 is the most balanced vaccine candidate, showing favourable safety, strong TLR4 interaction, broad epitope coverage, and robust immune memory, making it the top priority for development. *Na*-ASP-2 has strong immunogenicity but safety concerns that require engineering to reduce IgE-related risks while retaining efficacy. *Na*-GST-1 needs further experimental validation to clarify whether its low response reflects true immunogenic limitations or immune tolerance.

### Limitations

This study relies exclusively on *in silico* computational approaches to assess the immunogenicity and safety of the vaccine candidates. Although these methods provide valuable predictive insights, their findings require experimental validation in biological systems. Key limitations include:

1. The 3D structures were computationally predicted rather than experimentally determined, introducing structural uncertainties that may influence epitope mapping and molecular docking accuracy
2. Molecular docking and normal-mode analysis provide structural predictions but do not fully capture binding kinetics, antigen processing, or MHC presentation dynamics that are critical for actual immune activation.
3. The study does not account for immune responses in hookworm-exposed populations, where pre-existing sensitization, particularly to *Na*-ASP-2, may significantly influence vaccine safety and efficacy.
4. Population coverage based on HLA predictions is approximate. Coverage depends on predicted MHC binding and available allele frequency data; actual presentation may differ, and real-world immunogenicity can vary among individuals and regions.

## Conclusion

This study employed an integrated immunoinformatics pipeline to retrospectively evaluate the safety and immunogenicity of three hookworm vaccine candidates, *Na*-APR-1, *Na*-GST-1, and *Na*-ASP-2, against *Necator americanus*. The computational analysis correctly flagged *Na*-ASP-2 as toxigenic and allergenic, consistent with its clinical failure, while *Na*-APR-1 showed the most favourable safety profile. Although *Na*-GST-1 exhibited conflicting allergenicity predictions, all three candidates contained diverse immune-stimulating epitopes, with *Na*-APR-1 showing the broadest epitope coverage and *Na*-ASP-2 the highest predicted antigenicity. All three antigens demonstrated more favourable TLR4 binding energies compared to the positive control agonist, with *Na*-GST-1 and *Na*-APR-1 exhibiting the strongest predicted affinities and stable antigen- TLR4 complex dynamics. Immune simulation revealed *Na*-APR-1 and *Na*-ASP-2 elicited robust, memory-driven humoral and cellular responses, while *Na*-GST-1 showed markedly attenuated simulated immunogenicity, a finding attributed to model limitations rather than true absence of immunogenic potential, given established clinical evidence. Collectively, these findings position *Na*-APR-1 as the most balanced and clinically promising candidate, advocate for targeted safety engineering of *Na*-ASP-2, and highlight the value of integrated computational validation as a complementary framework for guiding rational hookworm vaccine development.

## Supporting information

**S1 Table 1.** Characteristics of predicted conformational epitopes.

**S1 Table 2.** Some predicted cytokine-inducing epitopes.

**S1 Table 3.** Binding affinity of the vaccine candidates and agonist.

**S1 Fig 1.** Molecular docking protein-protein interaction pocket prediction and of the vaccine candidates with the TLR4 receptor was conducted

**S1 Fig 2.** Protein-protein interaction pocket prediction

**S1 Fig.3** Population coverage analysis of predicted T-cell epitopes across the global population is presented as follows

**S1 Fig.4** Population coverage analysis of predicted T-cell epitopes across the global population is presented as follows

**S1 Fig 5**. Predicted intrinsic disorder profiles of the *N. americanus* vaccine candidates, *Na*-APR- 1, *Na*-ASP-2, and *Na*-GST-1.

## Acknowledgments

The authors acknowledge all the open-source and globally available databases utilized for data mining.

## Author Contributions

**Conceptualization:** Y.A.S.T., and R.A.S.; **Methodology**, Y.A.S.T., R.A.S., G.T.N., A.B.A., K.P.N., N.G.N., T.M.E.; **Software**, Y.A.S.T., R.A.S., A.B.A., M.N.M., T.B.M., N.E.Y., B.N.Y., D.M.B., K.Y.G., B.T.T., C.M.S.; **Formal analysis**, Y.A.S.T., R.A.S., T.M.E., G.T.N., A.B.A., K.P.N., C.R.A., M.S.L., K.Y.G., C.M.S., B.T.T., M.N.M., N.G.N., S.M.G.; **Writing-Original Draft Preparation**, Y.A.S.T., R.A.S., G.T.N., M.S.L., B.N.Y., J.E.E., **Writing-Review and Editing**, All authors.; **Visualization**, Y.A.S.T., R.A.S., T.M.E., and G.T.N.; and **Supervision**, S.M.G., and R.A.S.

## Institutional review board statement

Not applicable.

## Informed consent statement

Not applicable.

